# High-throughput biochemical profiling reveals Cas9 off-target binding and unbinding heterogeneity

**DOI:** 10.1101/059782

**Authors:** Evan A. Boyle, Johan O. L. Andreasson, Lauren M. Chircus, Samuel H. Sternberg, Michelle J. Wu, Chantal K. Guegler, Jennifer A. Doudna, William J. Greenleaf

**Author notes:** Equal Contribution (Evan A. Boyle, Johan O. L. Andreasson, Lauren M. Chircus).

## ** Introduction

The bacterial adaptive immune system CRISPR-Cas9 has been appropriated as a versatile tool for editing genomes, controlling gene expression, and visualizing genetic loci^1^. To analyze Cas9’s ability to bind DNA rapidly and specifically, we measured the kinetics of catalytically dead Cas9 (dCas9) interactions with a library of potential binding partners. Using a massively parallel assay of protein-DNA interactions derived from a high-throughput sequencing flow cell (HiTS-FLIP)^2^ and building on the established importance of protospacer adjacent motif (PAM) and seed recognition^3–5^, we identify two PAM-distal regions, proximal and distal to the seed region, with distinct behaviors: multiple mismatches in the seed-proximal region work in a highly synergistic manner to reduce Cas9 association whereas seemingly tolerated mismatches in the distal region precipitate comparatively rapid dissociation of Cas9. Together, these observations support a model for Cas9 specificity wherein gRNA-DNA mismatches at distinct domains of PAM-distal bases modulate different biophysical parameters of association and dissociation, opening the possibility of kinetic and thermodynamic tuning of the Cas9-DNA interaction and quantitative prediction of off-target binding behaviors.

## ** Main body

CRISPR-associated protein 9 (Cas9) is programmed to bind its target DNA by a guide RNA (gRNA) that, once loaded, forms a ribonucleoprotein (RNP) complex. The *Streptococcus pyogenes* CRISPR system, the most extensively used system to date, targets a 23-bp DNA sequence containing 1) an ‘NGG’ PAM element in the non-template strand of the target DNA and 2) a 20-bp sequence upstream of the PAM bearing complementarity to the gRNA. Genome engineering applications leverage the nuclease activity of the Cas9 RNP, but catalytically inactive Cas9 (dCas9) has proven valuable on its own by enabling the creation of customizable and programmable DNA binding elements that can activate and repress gene expression with high precision^1^.

The biophysical underpinnings of the Cas9 target search have been investigated both by directed biochemical assays ^6,7^ and through measurements of off-target Cas9 activity^5,8–12^. From these studies, it is clear that Cas9 proceeds through a series of steps starting with PAM recognition, followed by DNA melting, RNA strand invasion, and heteroduplex formation dependent on complementarity with a 5–10-bp seed. Structural data have further suggested that conformational changes in the HNH domain reposition catalytic residues and permit allosteric regulation of the RuvC domain. This conformational gating ensures that cleavage occurs only in the context of substantial homology between gRNA and target^13,14^.

Though Cas9 specificity is mediated through multiple mechanisms, the specificity of binding to DNA substrate is crucial for all potential applications of Cas9’s RNA-programmable targeting. ChIP-seq experiments have indicated that Cas9 stably binds sequences even with multiple mismatches at the most PAM-distal bases^10,11,15^; however, ChIP-seq data does not perfectly recapitulate the Cas9 target search, and no scalable approach exists for exhaustive profiling of off-target binding *in vitro.* Determining the full extent to which sequence controls Cas9 binding specificity, including kinetic rate constants, is thus a priority for creating a predictive and quantitative understanding Cas9 behavior at both on-and off-target sites.

To investigate the sequence determinants of Cas9 binding, we performed a comprehensive survey of dCas9 off-target binding potential. We generated a library of mutant targets on a massively parallel array and assessed binding of fluorescently labeled dCas9-sgRNA complexes in real time (Online Methods). We chose a well-characterized 20-bp phage lambda DNA target sequence^6,14^ and used mixed base oligonucleotide synthesis to construct a library of modified targets with maximal coverage of double substitutions. Illumina sequencing adapters were incorporated to flank these target sequences (Fig. 1a), permitting cluster generation and sequencing on an Illumina flow cell. Upon completion of sequencing, the GAIIx flow cell comprised a two-dimensional array of clonal DNA clusters with the template strand tethered to the surface of the flow cell. Each cluster of identical potential DNA binding partners contained anywhere from zero to twenty substitutions from the 23-nucleotide on-target sequence.

After using the high-throughput sequencing data to define the spatial coordinates of sequence clusters in the library, the flow cell was placed into a modified GAIIx instrument^16^ for biochemical profiling. Programmed RNP complexes were introduced into the flow cell at either 1 nM or 10 nM concentration (Supplementary Table 1) and left to incubate 12 hours overnight to approach saturation. Following this incubation, dCas9 was displaced with dCas9-free buffer, either with or without a high concentration of unlabeled double-stranded target DNA to minimize dCas9 rebinding to DNA targets following dissociation. During association and dissociation experiments, image series were collected in real time across all 120 tiles of the flow cell lane (Fig. 1b). For each experiment, initial apparent on-rates and off-rates were calculated by fitting fluorescence values for each off-target, corresponding to observed pre-steady state on-rates and initial observed off-rates. (Fig. 1c,d, Online Methods).

Apparent on-rates were obtained for 84,554 sequences, including all single mutants, 99% of all possible double mutants, and 59% of all possible triple mutants, as well as 64,594 higher order mutants (Fig. 1e). Datasets (1 and 10 nM) were merged to evaluate all target apparent on-rates jointly (see Online methods, Supplementary Fig. 1). Single mutants were generally measured across >1,000 clusters. Sequences with four or more mismatches were typically measured across twenty or fewer clusters due to the combinatorics of introducing higher order mutants (Fig. 1f). Across single and double mutants, dCas9 on-rates were highly reproducible (*R*^2^ = 0.962) (Fig. 2a). Amongst all potential binding targets, reproducibility was slightly reduced due to lower cluster counts per sequence (*R*^2^ = 0.856, Supplementary Fig. 2).

Examination of apparent on-rates for single mutants in the guide RNA complementary nucleotides (positions −20 to −1) confirmed that substitutions in a ~7-bp seed region (positions −1 through −7) were largely responsible for reducing on-rates relative to the wild type sequence (Fig. 2b). Substitutions outside this seed region were more muted in their affect on association rates. Although seed region mismatches posed greater barriers to dCas9 binding than PAM-distal mismatches (positions −8 to −20), we found that both position and base identity of a mismatch were required to determine the initial apparent on-rates of off-target binding (Fig. 2b). The identity of the degenerate base in the NGG PAM (position +1) had no detectable effect on apparent on-rate kinetics (KS test *p* > 0.05), consistent with prior observations^9, 12, 17^. Because others have observed that off-target DNA sequences can exhibit higher Cas9 cleavage efficiencies than their on-target site^5,18,19^, we wondered whether such off-targets existed in our data. At a 1% Benjamini-Hochberg FDR significance threshold, we found no evidence for off-target sequences exhibiting apparent on-rates exceeding that of the on-target site.

As expected, dCas9 bound targets that were mutated at the GG dinucleotide of the PAM (positions +2 and +3) at undetectable or nearly undetectable rates. For these PAM-mutated targets, apparent on-rates were largely not influenced by sequence complementarity between guide RNA and off-target DNA; however, nearly all constructs in the flow cell contained at least one GG dinucleotide, often present in the barcode or introduced in the target sequence itself. Thus, we infer that the slowest association rates we observed, approximately fifteen-fold below the rate for the on-target sequence, represent PAM-scanning behavior. In support of this, we find that PAM mutant targets are less likely to show detectable levels of binding than mismatched targets with intact PAMs (11% versus 47%). Furthermore, PAM mutants that do exhibit detectable binding contain a greater number of novel GG dinucleotides on average than PAM mutants that do not bind, both on the non-template strand (0.71 vs 0.54 novel GGs, *p* < 1·10^−280^, Wilcoxon rank sums test) and on the template strand (0.61 vs. 0.48, *p* < 1·10^−280^). In contrast to ChIP-seq experiments that yielded no recurring non-canonical PAMs, but consistent with other literature^5,20,21^, we observe that mutations of the NGG PAM to NGA or NAG permitted more accumulation of Cas9 signal over time relative to other PAM mutants. These non-canonical PAM targets possessed initial apparent on-rates comparable to all other PAM mutants, which may be due to low resolution in quantified association rates at low levels of binding or differences in thermodynamic rather than kinetic properties.

Next we asked if the effect of combining multiple substitutions in one off-target could be predicted from the energy barriers to binding of the constitutive single mutants – that is, whether effective energy barriers governing on-rates were additive. For this analysis, PAM and seed mutants were not assessed, as canonical binding was largely abrogated and any observed binding was presumed to reflect PAM-scanning behavior independent of the number of mismatches. In the PAM-distal region we found two distinct milieus of negative epistasis (non-additivity) (Fig. 2c). For many PAM-distal bases (positions −12 to −20), double mutants exhibited slightly lower on-rates than expected from the single mismatch contributions. The four bases adjacent to the seed (−8 to −11), however, showed much larger decreases in on-rates when two mismatches were present. Curiously, although the three terminal PAM-distal bases (−18 to −20) are considered dispensable for binding^22^, and single mismatches in this region produced little change in on-rate, we found that the presence of a second mismatch in the four bases adjacent to the seed (−8 to −11) greatly sensitized dCas9 to mismatches in the terminal nucleotides (Fig. 2c, Supplementary Fig. 3). This phenomenon was also evident across higher order (>2) mutants (Fig. 2d). This distinct behavior prompted us to divide the PAM-distal region (−8 to −20) into two sections: a single-mutation tolerant zone proximal to the seed (−8 to −11) and a multiple-mutation tolerant zone more distally (−12 to −20). These observations provide experimental support for a model of gRNA strand invasion that posits a critical role for PAM-distal bases in stabilizing Cas9 R-loops^23^ and highlight the difficulty of predicting Cas9 off-target activity at sites with multiple mismatches.

We next turned our attention to dCas9 occupancy following the 12-hour overnight incubation. Prior work has established that Cas9’s interactions with on-target sites are very stable^24^ and that Cas9 interrogation of PAMs without seed homology is extremely short-lived^25^. More poorly understood is stability of binding of off-targets with a small number of substitutions from the on-target sequence. We confirmed that most, but not all, double substitutions in the seed or PAM regions abrogated dCas9 occupancy (Supplementary Fig. 4). In contrast, the vast majority of single substitutions achieved high levels of occupancy, even in cases with very slow on-rates. Amongst other single and double substitutions, there was considerable variation in the fraction of DNA ultimately bound by dCas9, suggesting that, for some sequences, association and dissociation of dCas9 were at equilibrium for an intermediate level of occupancy.

Limited experimental data^23^ and modeling efforts^26^ have suggested a role for variation in dCas9 off-rates mediated by target mismatches, but to our knowledge there has been no systematic characterization of how specific target mismatches effect changes in either Cas9 or dCas9 off-rates. To characterize the Cas9 binding process more completely, we thus directly measured the initial observed off-rates of dCas9 from off-target DNA.

Surprisingly, we found high variation in apparent initial off-rates across dCas9 off-target sequences (Fig. 3a), in addition to the variation we found in the level of saturated signal. In contrast to dCas9 on-rates, which were primarily determined by complementarity in the seed region, apparent dCas9 off-rates appear to be almost exclusively modulated by the PAM-distal region, more specifically a region spanning positions −8 to −17, which we define as the reversibility-determining region, or RDR. Though dissociation was immeasurably slow for the on-target sequence, we found that a single mismatch in the PAM-distal region (−16G) induced near complete dissociation of dCas9 in less than an hour. From this data, it appears that, when gRNA strand invasion bypasses seed mismatches in off-target DNA, a highly stable complex is formed via favorable PAM-distal base-pairing. In contrast, R-loop formation over RDR mismatches may lead to dissociation of dCas9 on the time scale of minutes to hours, even when the seed region is an exact match. This variation in off-rates does not appear to be an artifact of differences in on-rates (Supplementary Fig. 5), and the −16G construct along with several other test sequences were validated for association and dissociation characteristics by filter binding assays (Supplementary Fig. 6). Adding unlabeled competitor DNA resulted in systematically higher dCas9 off-rates, but these data were strongly correlated with passive flow experiments (*R*^2^ = 0.752, Supplementary Fig. 7).

Our high-throughput profiling also showed that multiple mismatches in the PAM-distal region, especially the most distal bases of the RDR, trigger faster dissociation than that of the single mismatches alone (Fig. 3b). This observation prompted us to further subdivide the RDR into proximal (−8 to −11) and distal (−12 to −17) regions. It is worth noting that the differences in degree of epistasis divided the target DNA into precisely the same partitions: with the proximal RDR exhibiting much more extreme epistasis than the distal RDR. Though gRNA strand invasion has been envisioned as a single continuous process^24^, the data we have shown suggest that the gRNA:DNA heteroduplex can be thought of as a series of discrete regions that operate sequentially to determine whether and how stable R-loop formation occurs.

We next sought to relate our observed on-and off-rates to Cas9 off-target cleavage as predicted by an *in vivo*-derived model of cleavage efficiency^27^. Though we profiled binding of only one sgRNA, we could confirm that the on-rates calculated for dCas9 off-targets are predictive of modeled Cas9 off-target cleavage probabilities (*R*^2^ = 0.539, p = 2.3 × 10^−12^; Supplementary Fig. 8). Finally, we observe that within the RDR, predicted off-target cleavage activity is similarly correlated with off-target off-rate (spearman ρ = −0.35) as with off-target on-rate (spearman ρ = 0.33, Supplementary Fig. 8), and amongst all off-targets, predictions of model-derived cleavage efficiency are significantly improved (*p* = 8.5·10^−4^, Online Methods) when estimated using on-and off-rates (*R*^2^ = 0.559) as opposed to on-rates alone (*R*^2^ = 0.397).

Drawing from these observations, we propose a mechanism for both Cas9 binding and dissociation (Fig. 3c). PAM mutations act to rapidly release diffusing Cas9 molecules post-collision, whereas seed mutations impair target melting. When DNA melting and seed hybridization is accomplished in spite of seed mismatches, heteroduplex formation will continue to completion, and Cas9 will thus remain stably bound. Mismatches in the nucleotides adjacent to the proximal RDR modulate the energy barrier to dissociation such that heteroduplexes are reversed on a shorter timescale, especially when multiple PAM-distal mismatches are present. Finally, mismatches in the terminal nucleotides of gRNA-template pairing have little effect in isolation, but still destabilize the full heteroduplex and sensitize Cas9 to any additional mismatches in PAM-distal bases.

Because this assay measures only whether dCas9 is associated with DNA template, we cannot definitively link Cas9 structural features to individual steps of the coordinated multistep processes of binding and unbinding. Also, the apparent off-rates we measured are conditioned on high dCas9 occupancy following the association experiment. Thus, rapid off-rates of dCas9 from PAM and seed mutant off-targets are expected to manifest as low apparent on-rates rather than high apparent off-rates in our data.

Our results reveal the complex effects of combinatorial DNA sequence perturbations on the binding behavior of Cas9 and motivate further study into the role of Cas9 dissociation in increasing the specificity of both Cas9-mediated binding and cleavage. We identify observable dissociation kinetics as a novel functional consequence of PAM-distal mismatches in a set of nucleotides we term the reversibility-determining region, with especially acute consequences in the distal RDR. Modulation of Cas9 off-rate kinetics represents a promising area for novel application of CRISPR-Cas systems to tune thermodynamic and kinetic parameters for maximal efficiency or specificity, and may already underlie alternate genome editing approaches including truncated guide RNAs and modified Cas9 proteins^28–30^. We anticipate that these and other similar methodologies will provide a new avenue of molecular characterization – high-throughput biochemical profiling – which will facilitate functional dissection of novel nucleic acid-binding molecules in addition to other members of the CRISPR/Cas family of enzymes at unprecedented scale.

**Figure 1.**
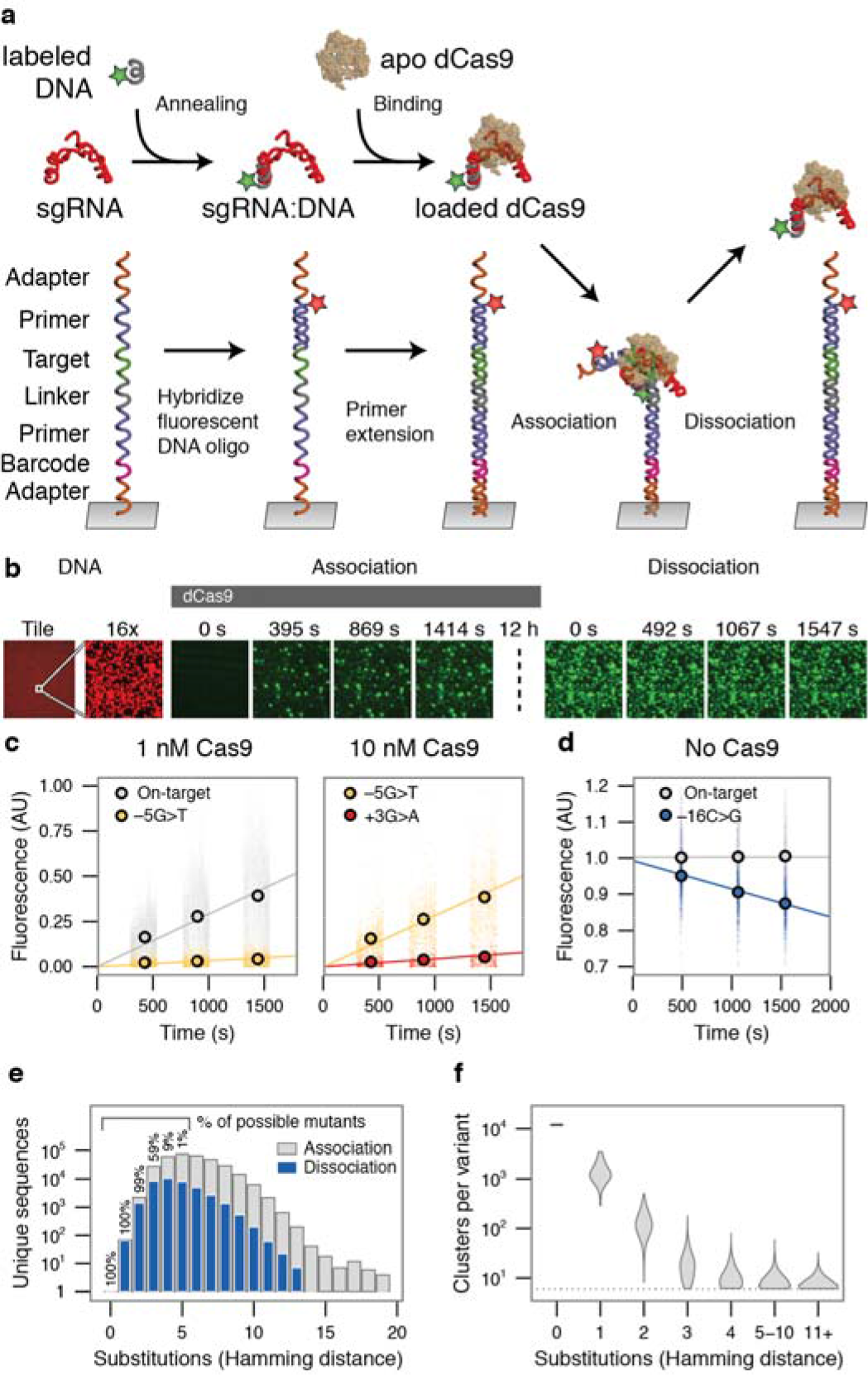
Quantifying dCas9 binding behavior on a massively parallel array. a) Experimental procedure for high-throughput biochemical profiling. A fluorescent DNA oligo hybridized to the dCas9 sgRNA was loaded into the apo-dCas9. In parallel, an Illumina sequencing-compatible DNA construct was both labeled and made doublestranded by extending a second fluorescent oligo. dCas9 was flowed into the chamber, allowing association with double-stranded DNA. A dissociation experiment was then performed by quantifying the decrease in dCas9 signal upon dilution or chase. b) Example images taken in two channels on the array, DNA in red and dCas9 signal in green. A 12-hour incubation, meant to saturate the clusters with dCas9, separates association from dissociation experiments (dotted line). For most clusters, signal accumulated in the on-rate experiment largely remains throughout the dissociation. c,d) Examples of c) association and d) dissociation lines fit to different targets The +1 base refers to the first base of the PAM, −1 to the most PAM-proximal base, and −20 to the most PAM-distal base. e) The total number (y-axis) and percentage (in text) of possible targets profiled for each number of substitutions from the on-target site. Only a fraction of sequences with quantified on-rates are profiled for off-rates (blue) with high confidence. f) Clusters per variant for targets with the given number of substitutions.

**Figure 2.**
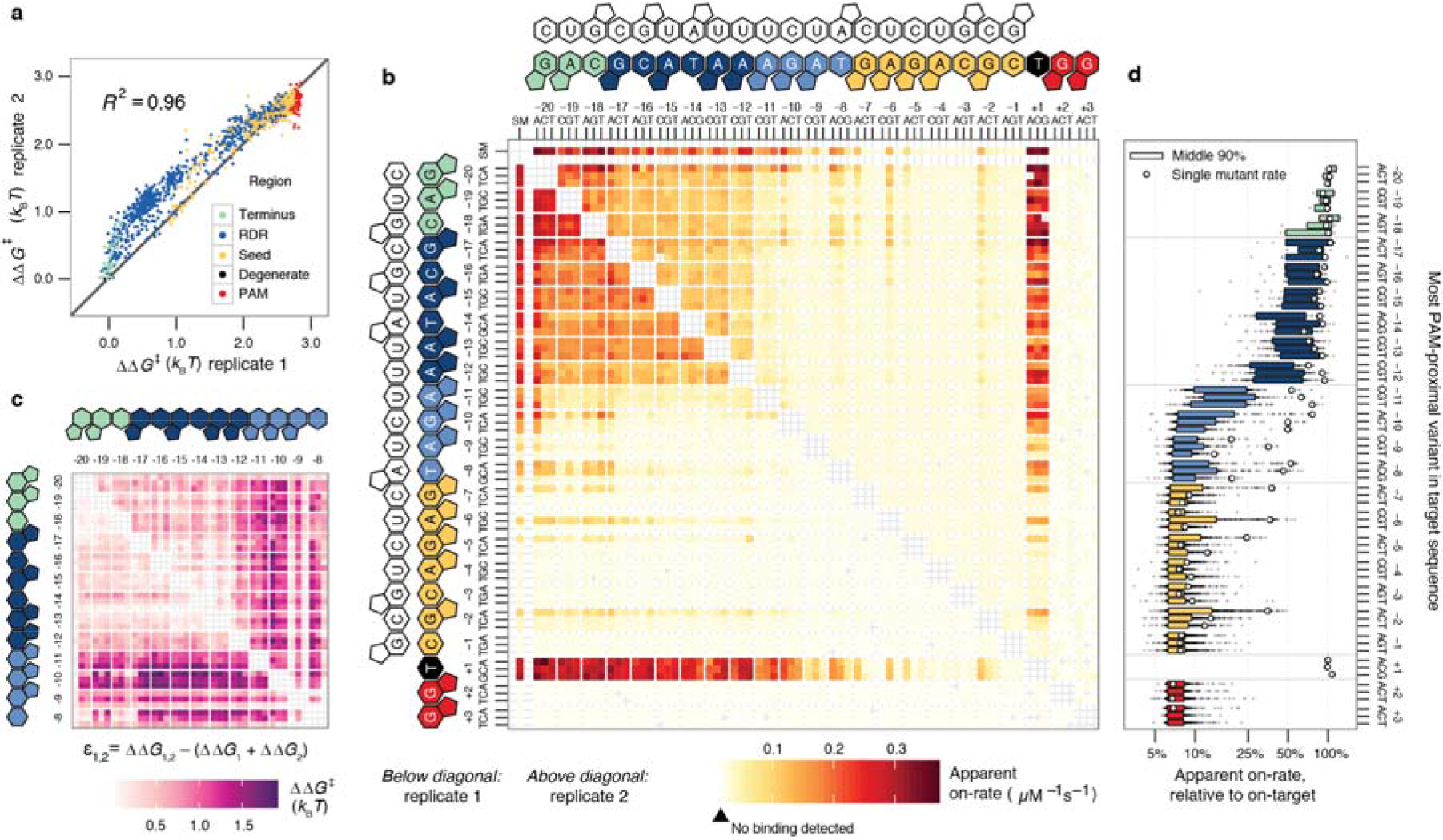
Deep profiling of dCas9 observed initial on-rates across a range of potential off-target sequences. **a)** dCas9 relative energy barrier reproducibility for single and double mutants across replicates, calculated relative to the on-target DNA. Points are colored by the more PAM-proximal mutation position, excepting the degenerate base of the ‘NGG’ PAM. **b)** Apparent on-rates for all single mutants (the series of tiles ‘SM’, horizontal and vertical) and double mutants (all other tiles) across both replicates, shown above and below the diagonal. Heat reflects higher on-rate for off-target sequences with the substitutions indicated on the x-and y-axes. Targets with at least 6 clusters but with no detectable binding were colored the minimum quantified rate. Double mutant cells lacking 6 clusters are left unfilled. **c)** Epistasis in energy barriers for double mutants for the PAM-distal nucleotides. Nearly all pairs of mismatches appear to lower on-rates more than expected by single mismatch estimates. The seed region shows little variation in on-rates, likely due to PAM scanning behavior, and is not shown. **d)** Distribution of higher order (>2) mutant on-rates summarized by their most PAM-proximal mutation (degenerate base excluded).

**Figure 3.**
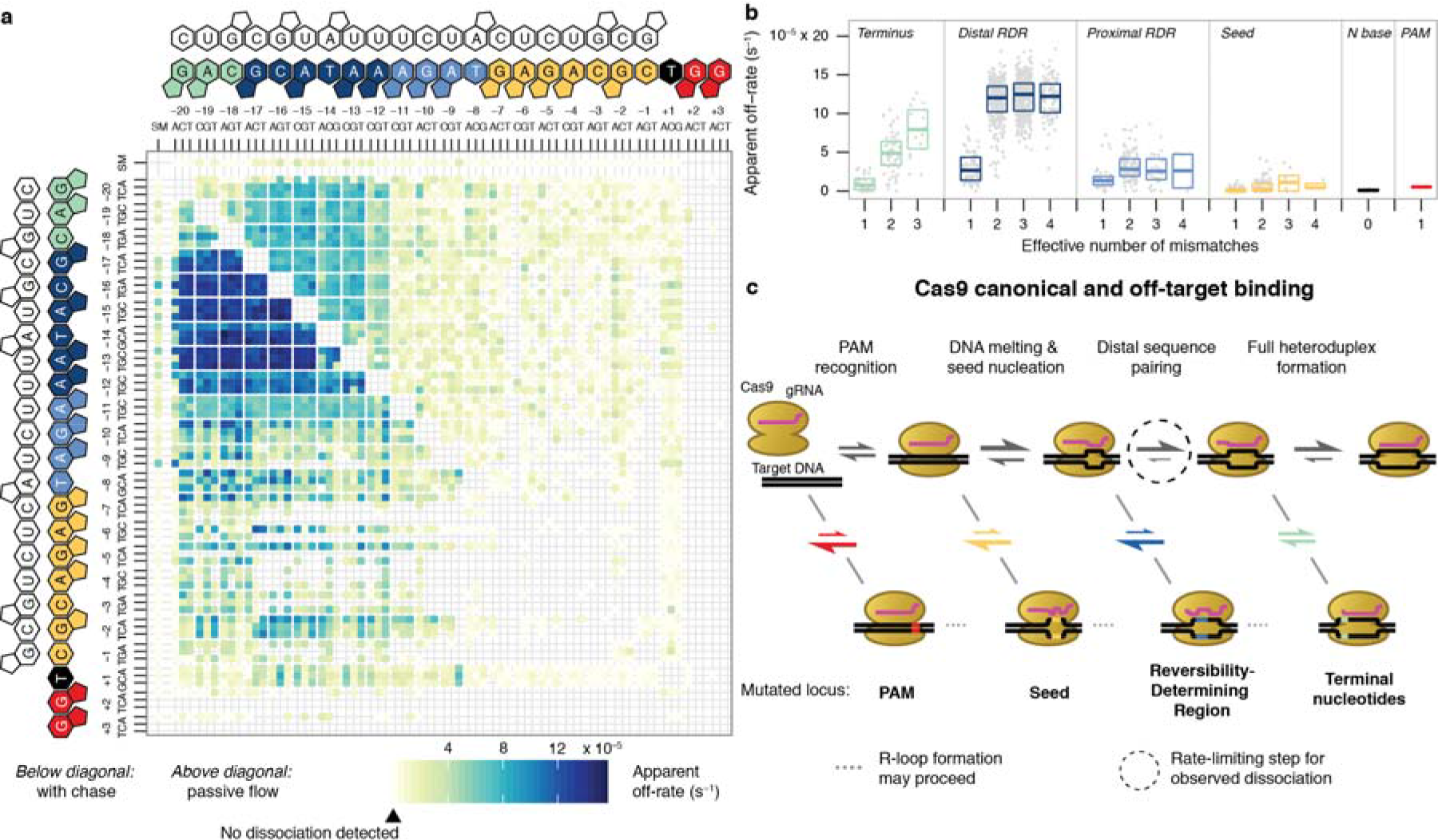
Variation in dCas9 off-rates suggests a model of Cas9 binding behavior. a) dCas9 apparent off-rates for single and double mutants, (see Fig. 2b). Apparent off-rates are systematically higher in the presence of an unlabeled competitor dsDNA that prevents rebinding (below diagonal) than without (above diagonal). **b)** Effect of multiple mismatches on off-rates partitioned by region of the target. **c)** Model diagram for R-loop formation in different off-target contexts. Rates for protospacer mismatches are color-coded by off-target partition as in panel **a**. PAM and PAM-proximal mismatches affect early steps in the Cas9 target identification procedure, whereas distal mismatches influence later steps. Off-rates in panel **a** are likely products of the kinetics of unwinding of the R-loop across the RDR. Off-rates associated with transient binding, as with most PAM mutants that fail to form R-loop structures, do not appreciably bind and thus are not captured in the dissociation experiment.

## ** Online Methods

### * dCas9 and sgRNA preparation

dCas9 (the catalytically dead D10A/H840A mutant) was purified as described^17^. The sgRNA (Supplementary Table 2, item 1) was *in vitro* transcribed from the BamHI cleavage product of pSHS 256 (benchling.com/s/zmUR5HNi/edit) using T7 polymerase. The 3′ end of the sgRNA was extended to permit annealing of the Cy3 probe. Both the 3′ extension and hybridized Cy3 probe were loaded into dCas9 and tested on on-target DNA to ensure no defect in Cas9 binding resulted.

### * Library design

The DNA library was constructed from two DNA oligos synthesized by the Stanford Protein and Nucleic Acid Facility. One DNA oligo contained Illumina P7 sequence, a 15 base randomer to serve as a unique molecular identifier barcode, and custom sequencing primer cspLC 01. The second DNA oligo contained Illumina P5 sequence, custom sequencing primer cspLC 02, and the doped lambda 1 dCas9-sgRNA target binding site. Degenerate bases were introduced using hand mixed bases with a 76% chance of incorporating the sgRNA on-target base and an 8% chance of incorporating each alternative base. The degeneracy was optimized with the intent to cover all double mutants.

### * Library assembly

Libraries were assembled by annealing two oligos and extending with Phusion HF polymerase (NEB) to generate double stranded library members. Excess single stranded oligos were digested with ExoI (NEB). The number of full-length molecules in each library was then quantified by qPCR, and the population of each library was bottlenecked to approximately 950,000 unique molecules, each of which could be identified by its unique 15-base barcode. This bottlenecked library was then amplified to 30 nM by PCR and quantified as previously described.^16^.

### * Sequencing

Elim Biopharmecuticals (Hayward, CA) performed single-end sequencing of the library on an Illumina GAIIx sequencer and generated 15,652,121 clusters on one lane of a flow cell. Sequencing consisted of a 15-cycle barcode read (cspLC 01) followed by a 27-cycle target read (cspLC 02). To ensure proper cluster assignment in subsequent kinetic profiling on the flow cell, clusters not passing filter were included in downstream image analyses.

To improve barcode mapping accuracy, the library was also sequenced on an Illumina HiSeq sequencer by Elim Biopharmecuticals (Hayward, CA). 2x50 paired-end sequencing was performed with the same barcode read primer (cspLC 01) and a paired-end target read primer (cspLC 03) to free the index read. This yielded 43,617,449 total reads for mapping barcode sequences to target sequences.

### * Barcode analysis

The sequencing data from both the GAIIx and HiSeq sequencing runs were combined in the following way: target reads from both runs were trimmed to the first 23 bases (the length of the doped target sequence), where GAIIx target reads were reverse complemented to share the orientation of the HiSeq target reads. The HiSeq barcode read was trimmed to the first 15 bases (the length of the barcode). Thus, each cluster could be summarized by cluster ID, barcode read, barcode read q-scores, target read, and target read q-scores.

To ensure high sequencing quality, a barcode-target dictionary was generated in a base-wise manner: for each position in the 23-bp target sequence, the most common base across the barcode’s clusters was called as the true base. After this, each barcode was annotated with (1) the consensus target sequence, (2) the number of clusters attributed to the barcode, (3) what proportion of target reads matched the consensus target sequence. Data from a barcode were removed from downstream analysis if they (1) had fewer than 4 clusters informing the mapping, (2) surpassed the mean number of clusters per barcode by 2 standard deviations, (3) represented a homopolymer, (4) aligned to the Illumina adapter sequence, or (5) had under 50% of contributing cluster target reads perfectly matching the consensus target read. The remaining barcodes were considered high quality and their consensus target sequences were considered the correct target sequence for that barcode.

### * Flow cell preparation

Following sequencing, the GAIIx flow cell was placed on a GAIIx sequencer modified as previously described^16^. The second strand of DNA generated during sequencing was stripped with formamide, and the residual fluorophores from sequencing were cleaved with cleavage buffer (100 mM TCEP, 100 mM Tris pH 8.0, 125 mM NaCl, and 0.05% Tween 20).

### * Association and dissociation experiments

The 3′ end of the sgRNA was labeled prior to loading onto dCas9 with a Cy3 labeled oligo (Supplementary Table 2, item 2) by incubating 4.95 μM sgRNA with 5 μM of the labeled oligo in hybridization buffer (20 mM Tric-HCl pH 7.5, 100 mM KCl, 5 mM MgCl_2_) for 5 minutes 95°C then slowly cooling to room temperature. For each experiment (1 nM and 10 nM dCas9), the specified concentration of dCas9 was incubated with 50 nM labeled sgRNA at 37°C for 25 minutes in binding buffer (20 mM Tric-HCl pH 7.5, 100 mM KCl, 5 mM MgCl_2_, 5% glycerol, 0.05 mg/mL heparin, 1 mM DTT, and 0.005% Tween 20) to load the sgRNA onto the dCas9. Labeled sgRNA was confirmed to yield no quantifiable association signal on the flow cell surface in the absence of dCas9, so excess sgRNA was used to ensure that all dCas9 molecules bound sgRNA. Each loaded dCas9-sgRNA preparation was placed on ice throughout the course of the experiment.

Prior to each association and dissociation experiment, any DNA hybridized to the DNA tethered on the flow cell surface in a previous experiment was removed with a 100 mM NaOH solution. Next, an Alexa-647-labeled oligo (Supplementary Table 2, item 3) was annealed to common sequence on the tethered DNA. dsDNA was generated by extending the annealed oligo with Klenow Fragment (3′ → 5′ exo-) (NEB) in buffer per manufacturers’ recommendations, 30 min at 37°C.

Following dsDNA generation, the loaded dCas9-sgRNA was perfused over the flow cell. The lane containing the dCas9 target library was imaged for approximately 25 minutes following the initial perfusion, 4 minutes per 120-tile image series plus pauses between series. dCas9-sgRNA was replenished at varying intervals, with no changes in observed on-rates as a result of this at the time-scales investigated.

After approximately 12 hours of dCas9 perfusion, another image series was taken to confirm saturation of binding of dCas9 on the DNA clusters. Following this series we performed the dissociation experiments. In one experiment, 250 nM unlabeled competitor on-target dsDNA in binding buffer was introduced into the flow cell to replace the dCas9 solution. The on-target acts as a competitor to prevent soluble dCas9-sgRNA from rebinding clusters on the surface after dissociation. In the passive flow experiment, binding buffer without target DNA was used instead. We suspect that the high local concentration of DNA in flow cell clusters, the possibility of protein aggregation over time, and a nontrivial rate of photobleaching per image (estimated to be ~2% from consecutive imaging) could confound results over time. Therefore, we limited the time scale of each experiments to three informative image series, which should be relatively unimpaired by these biases.

### * Data Processing

Raw images were processed using software previously described^16^. Time stamps were extracted from image file metadata to assign the exact time the data were recorded. The time for the Illumina GAIIx to perfuse new solution over the flow cell and take the next image was found to be consistent across runs and enabled determination of how long dCas9 participated in binding prior to the first image taken.

To account for systematic differences in focus, cluster formation efficiency and illumination across tiles, fluorescence values were normalized per tile for on-rates by dividing them by the median quantified fluorescence amongst the on-target clusters in the post-saturation images collected 12 h after the initial perfusion. High background clusters were filtered out if their first quantified value exceeded 2% of the saturated signal for on-target binding. GG dinucleotides in oligo library barcodes and Illumina adapters as well as non-specific deposition could confound studies of PAM recognition and very low-level binding, so off-targets were filtered out if they failed to reach 2% of the saturated signal for the majority of their corresponding clusters. Initial on-rates were calculated by performing linear regression on the quantified fluorescence values across all clusters, constraining the fit to go through the origin. To permit joint analysis of 1 and 10 nM datasets, linear regression was performed on target sequences quantified for both concentrations, and 10 nM slopes absent in the 1nM dataset due to the limits of detection were inferred from the fit line (Supplementary Fig. 1).

Prior to calculating off-rates, each cluster’s quantified fluorescence values were further normalized. First, data at each time point was normalized by the on-target median fluorescence at that time. This accounts for additional variation in signal assuming that dissociation of dCas9 from its canonical target is negligible in the first 30 minutes and in practice amounts to a small adjustment. Next, each cluster's intensities were normalized by the first data point in the dissociation series such that corrected values represented the proportional decrease in fluorescence signal. DNA sequences with a median intensity for the first data point below 30% that of the on-target signal were excluded to filter out noisy fits.

Initial off-rates were also calculated by linear regression but without constraining the intercept. For both on-and off-rates, standard errors and confidence intervals were calculated by bootstrapping the clusters used in linear regression 100 times.

For comparisons of on-and off-rates to predicted cleavage efficiency, the “cfd-score-calculator.py” script was used to predict cleavage efficiency for each singly substituted target paired with the on-target guide RNA sequence. These predicted cleavage efficiencies were in turn used as labels to be predicted by dCas9 kinetic rates. Pearson correlations were calculated using out-of-bag predictions from the randomForest package in R with default parameters trained on on-rates and measured or permuted off-rates as covariates. Significance of inclusion of off-rates was assessed by assigning a standard error to estimates from 500 permutations of off-rates and calculating the probability of observing the measured pearson correlation from the standard normal distribution.

### * Filter binding experiments

DNA targets identical in sequence to the flow cell clusters were selected, using the most common barcode for each off-target. Six targets (on-target, −16G, −16T, −13C, −5T, and +3A) were ordered as gBlocks from IDT (Supplementary Table 1, items 4-9), amplified by PCR, and gel purified. The dsDNA was then 5′ radiolabeled by incubating by incubating 150 nM dsDNA, 1× T4 PNK (NEB), 1× PNK buffer (NEB) and 1 μM [*γ*- 32P]-ATP (PerkinElmer) for 30 min at 37°C followed by purification with a nucleotide removal kit (Qiagen). The sgRNA was hybridized to a labeled DNA oligo, as decribed above, and loaded onto the dCas9 by incubating 100 nM dCas9 and 125 nM sgRNA at 37°C for 25 min, and then at < 4°C.

Association rates were measured by incubating radiolabeled DNA targets with loaded dCas9 for different durations in a binding buffer identical to the flow cell experiments (see above). The total volume was 30 μL and the concentrations were either 10 nM dCas9 and < 240 pM DNA, or 1 nM dCas9 and < 150 pM DNA. For dissociation measurements, 10 nM dCas9 and < 240 pM target DNA were incubated for 2 h (on-target, −16G, −16T, −13C, +3A) or 5 h (−5T) followed by the addition of 6 μL non-radiolabeled cold competitor DNA in binding buffer (final conc. 83 nM). The quenching step lasted for different durations and both association and dissociation experiments were timed such that all conditions finished at nearly the same time. The samples were then applied to a 96-well BioDot^®^ microfiltration blotting apparatus under low vacuum, passing through a nitrocellulose membrane (Amersham Hybond ECL, GE Healthcare Life Sciences), a nylon membrane (Biodyne B, 0.45 μM, Thermo Scientific) and a filter paper (GE Healthcare Life Sciences) that were all pre-equilibrated with binding buffer. The membranes were allowed to dry, transferred to a phosphor screen overnight, and then measured on a Typhoon imager (GE Healthcare Life Sciences). Images were quantified in TotalLab Quant v12.2 and the dCas9-bound DNA fraction was calculated as signal from the nitrocellulose membrane divided by the total signal from the both the nitrocellulose and nylon membranes.

### * EMSA experiments

Cy5-labeled DNA targets were generated by PCR. To measure association rates, DNA was incubated with loaded dCas9 for varying times followed by a quench step with a high concentration of unlabeled competitor on-target DNA. Sequences and binding buffer were identical to the filter binding experiments. Concentrations were 100 pM DNA, 1 nM dCas9 and 80 nM competitor for the on-target target and 400 pM DNA, 10 nM dCas9, and 100 nM competitor for the −5T off-target. Following the quench, samples were resolved by gel electrophoresis on a 10% native polyacrylamide gel (Mini-PROTEAN, Bio-Rad) in TBE running buffer (Bio-Rad) for 30–60 min at 120 V at 4°C. Gels were imaged on a Typhoon imager (GE Healthcare Life Sciences) and quantified in TotalLab Quant v12.2.

## ** Acknowledgements

This work was supported by grants from the Beckman Foundation and from National Institutes of Health numbers 5R01GM111990 and 1P50HG00773501 to WJG. We thank members of the Greenleaf lab for feedback on data visualization. EAB acknowledges support from the NIH training grant 5T32HG000044-19. LMC acknowledges support from NIH training grant T32GM067586 and the National Science Foundation graduate research fellowship program. JAD is an Investigator of the Howard Hughes Medical Institute.

## ** Author contributions

LMC, SHS, JAD and WJG conceived of the study. EAB, JOLA, and LMC performed high-throughput biochemical profiling. EAB, JOLA, SHS, and CKG carried out EMSA validations. EAB and JOLA performed radioactive filter-binding assays. EAB, JOLA, LMC and MJW analyzed high-throughput data. EAB and JOLA wrote the text and generated figures. All authors reviewed and edited the manuscript.

## ** Competing financial interests

The authors declare no competing financial interests.

